# Evidence for a Single Rather than a Triple Dissociation in the Medial Temporal Lobe: an fMRI Recognition Memory Replication Study

**DOI:** 10.1101/2022.04.14.488400

**Authors:** Julie Van, Sam E. Nielsen, C. Brock Kirwan

## Abstract

Common research practices in neuroimaging studies using functional magnetic resonance imaging may produce outcomes that are difficult to replicate. Results that cannot be replicated have contributed to a replication crisis in psychology, neuroscience, and other disciplines over the years. Here we replicate two previous papers in which the authors present two analysis paths for a dataset in which participants underwent fMRI while performing a recognition memory test for old and new words. Both studies found activation in the medial temporal lobe including the hippocampus, with the first demonstrating a distinction in activation corresponding to true and perceived oldness of stimuli and the second demonstrating that activation reflects the subjective experience of the participant. We replicated the behavioral and MRI acquisition parameters reported in the two target articles (Daselaar, Fleck, & Cabeza, 2006; Daselaar, Fleck, Prince, et al., 2006) with N=53 participants. We focused fMRI analyses on regions of interest reported in the target articles examining fMRI activation for differences corresponding with true and perceived oldness and those associated with the subjective memory experiences of recollection, familiarity, and novelty. Comparisons between true and perceived oldness revealed main effects not only for true, but also perceived oldness along with significant interaction effects. We replicate the findings of recollection and familiarity signals in the hippocampus and medial temporal lobe cortex, respectively, but failed to replicate a novelty signal in the anterior medial temporal lobe. These results remained when we analyzed only correct trials, indicating that the effects were not due to selectively averaging correct and incorrect trials. Taken together, our findings demonstrate that activation in the hippocampus corresponds to the subjective experience associated with correct recognition memory retrieval.

**Highlights:**

- We replicated the methods of two influential papers using fMRI to examine recognition memory.
- The hippocampus consistently signals the subjective experience of remembering but not the objective old/new status of stimuli.
- Hippocampal response was consistent with recollection while the medial temporal lobe (MTL) cortex response was consistent with familiarity.
- We did not observe any regions in the MTL that signaled novelty, indicating recognition memory does not involve a triple dissociation of recollection, familiarity, and novelty in the MTL.

## 1. Introduction

Common research practices in neuroimaging studies using functional magnetic resonance imaging (fMRI) may produce outcomes that are difficult to replicate (Button et al., 2013; Simmons et al., 2011), leading to an ongoing crisis in psychology, neuroscience, and other disciplines in recent years (Shrout & Rodgers, 2018). Conflicting study conclusions may also impose interpretative errors that subsequently influence other relevant areas of research. Neuroimaging research generally involves lower samples of participants, consequently yielding low statistical power in studies and potentially inflating the rates of false positive outcomes and effect sizes (Button et al., 2013). Meta-analyses have identified over 1,000 research studies containing low samples (N’s = 12), with more recent studies in 2017-18 having included larger samples (N’s = 23-24; Szucs & Ioannidis, 2020). Although sample sizes in neuroimaging studies have increased over time, they remain relatively low, which in turn can lead to low replicability rates (Szucs & Ioannidis, 2020).

While more than half of surveyed scientists agree with the critical nature of the replication crisis, only less than 31% have adopted the view that failure to replicate may indicate false findings (Baker, 2016). Many, however, still retain their trust in published literature. However, when confronted with new information from replication results, most psychologists update their beliefs in effect sizes of previously-published studies (McDiarmid et al., 2021). To address this, our efforts to replicate focus on two highly influential works that informed other publications on recognition memory in the medial temporal lobe (MTL).

In two papers published in 2006, Daselaar and colleagues used identical data collection methods but two different analysis pipelines to investigate MTL responses to true vs. perceived oldness independent of subjective confidence in a recognition memory paradigm (Daselaar, Fleck, Prince, et al., 2006) and the influence of recognition confidence to examine the processes of recollection, familiarity, and novelty in the MTL (Daselaar, Fleck, & Cabeza, 2006). Recognition memory reflects the ability to acknowledge something as having been previously encountered. It is commonly thought to depend on recollection and familiarity processes, where recollection refers to memory retrieval accompanied by contextual details, and familiarity describes an impression that an item is old despite the absence of context that confirms this (Daselaar, Fleck, & Cabeza, 2006). Recollection is thought to be accompanied by the highest recognition memory confidence in an all-or-none manner while familiarity is more graded (Yonelinas, 2001). Novelty is similarly described as a graded process corresponding to the extent to which encountered information is discrepant from the expected information that an individual possesses (Skavronskaya et al., 2020).

Both papers found evidence of differential activation in the hippocampus and other MTL structures, indicating differentiation of responses between hippocampus and MTL cortex for processes supporting recognition memory performance. However, the different analysis paths lead to different conclusions about the role of the MTL and specifically the hippocampus in recognition memory. In the contrast of veridical old vs. new stimuli, regardless of behavioral response, the authors found increased MTL activation for true old stimuli (both hits and misses) compared to new stimuli (both false alarms and correct rejections) regardless of subjective experience as indicated by behavioral choices (Daselaar, Fleck, Prince, et al., 2006). On the other hand, when only considering subjective confidence ratings and not objective oldness of the stimulus, the authors found that the hippocampus selectively responded to the highest recognition memory confidence (often associated with the subjective experience of recollection) while different regions of the MTL cortex responded in a graded fashion to increasing familiarity or decreasing novelty (Daselaar, Fleck, & Cabeza, 2006). Thus, the same behavioral and fMRI paradigm yields evidence that the MTL both responds to the subjective experience of recollection as well as to the objective oldness of stimuli regardless of subjective experience, depending on the analysis path. Accordingly, we undertook a replication study matching behavioral procedures to the original papers and replicating analysis pathways with modern, open-source analysis tools in a much larger sample than the original study. We defined *a priori* regions of interest (ROIs) based on the original papers to perform parallel analyses examining MTL responses to A) objective vs. subjective oldness of test stimuli and B) subjective recognition confidence ratings corresponding to the experiences of recollection, familiarity, and novelty. Our hypotheses were that the same brain regions would demonstrate the same activation patterns observed in the original publications. Additionally, we performed exploratory voxel-wise analyses to test for activations more broadly within the MTL. Finally, in order to address a potential concern regarding selective averaging of correct and incorrect trials in the memory confidence analysis (Kirwan et al., 2009; Yu & Rugg, 2010), we performed the analyses based on recognition memory confidence ratings using only correct (hit and correct rejection) trials.

## 2. Methods

### 2.1 Participants

To determine the sample size, an *a priori* power analysis was done by extracting values from Figure 1 of (Daselaar, Fleck, Prince, et al., 2006) in the right hippocampus using a Web Plot Digitizer (Rohatgi, 2020), given that this region exhibited the smallest differences reported. Main effects were computed by averaging hits and misses, and CRs and FAs before converting SEM to SD and averaging again. The resulting values were entered into g*power to estimate an effect size of 0.46 (Faul et al., 2009). A sample of N = 54 was required to achieve a power of 0.95 with an error probability of 0.05 (t(1,53) = 1.67). To account for attrition, we enrolled N = 60 healthy young adults (38 females; mean age 21.3, SD = 2.25) from the local community.

**Figure 1.**
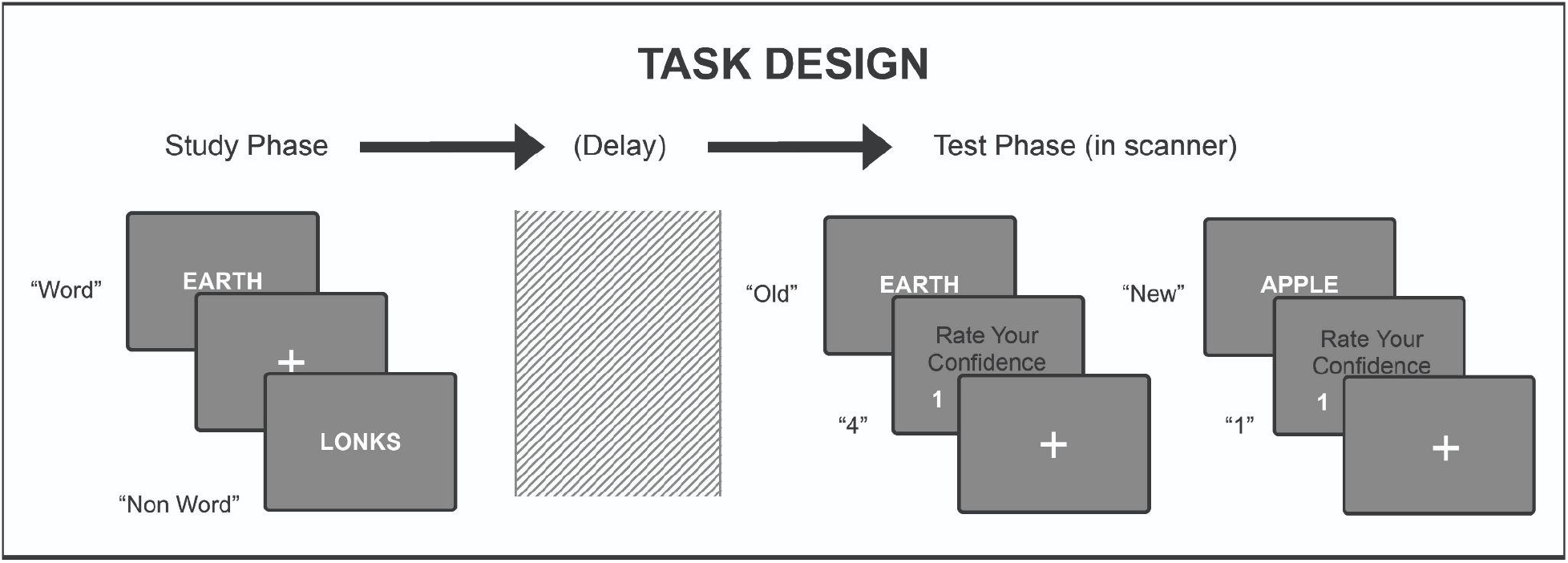
Study Protocol. Subjects were presented with a series of words and non-words at a computer during the study phase and asked to study the stimuli for later testing. After a subsequent delay, subjects were again presented with a series of words and non-words, some of which were included in the study phase and others that were entirely new. After indicating whether the stimulus was “old” or “new” they were prompted to rate their level of confidence.

Participants were recruited from the campus community and met MRI compliance screening requirements. Exclusion criteria included non-native English speakers, history of drug use, previous psychiatric or neurologic diagnosis, or contra-indications for MRI (e.g., ferromagnetic implant). We obtained written informed consent for all participants prior to study participation. Participants were compensated for participation with a choice of $20, course credit, or a 3D-printed ¼-scale model of their brains. The study was approved by the Brigham Young University Institutional Review Board. Data from n = 7 participants were excluded from MRI analyses for the following reasons: ineligibility (left-handedness, n = 1), failure to comply with study procedures (n = 2), excessive motion (n = 3), and equipment error (n = 1). Data collection and proposed analysis pipelines were registered to the Open Science Framework on 1 October 2020, prior to data collection (https://osf.io/ctvsw/registrations).

### 2.2 Stimulus Materials

The senior author of the original papers confirmed that the study materials used were no longer accessible (Cabeza, personal communication). Accordingly, we followed the protocols described in the original publications for the design of the paradigm. First, we composed a list of 240 five-letter words from the MRC psycholinguistic database (Wilson, 1988). All terms were of moderate frequency (ratings above 200), concreteness (ratings above 200), and imageability (ratings above 200). Stimuli were presented using PsychoPy3, a stimulus presentation and control package (Peirce et al., 2019). We generated a set of pseudo words by replacing vowels in the words of the stimulus list with different vowels using an R script. Stimuli and scripts are available on the OSF project page: https://osf.io/ctvsw/.

### 2.3 Procedure

Prior to MRI scanning, participants completed the study phase in which they were presented with a randomized list of 120 real English words and 80 pseudo words at a rate of 2000 ms per item (Figure 1). The instruction scripts that were read to participants can be found in the OSF project: https://osf.io/ctvsw/. A fixation cross was presented between these words for a random time interval varying between 0-5500 ms. Participants indicated whether the stimulus was a word or pseudo word using a number keypad. Participants were not informed at this time that their memory for the words would be tested. Following the completion of the study phase, researchers reviewed the completed screening forms with participants for MRI safety and participants were situated in the MRI scanner.

Once participants were situated in the MRI scanner, we obtained localizer, field map, and T1-weighted structural MRI scans (scan parameters below). Following these scans, we began the test phase of the experiment, approximately 30 minutes after the study phase as outlined in the original experimental protocols (Daselaar, Fleck, & Cabeza, 2006; Daselaar, Fleck, Prince, et al., 2006). The task paradigm was presented as four experimental runs lasting between 435-442 seconds. Participants saw an equal number of target stimuli (words shown during the study phase) and foil stimuli (novel words), with 60 words per run. Target and foil stimulus presentation order was randomized. Stimuli were presented for 3.4 seconds, and participants were instructed to make judgments indicating whether the word was presented on the study list while the stimulus was displayed. We then collected confidence ratings for those judgments between true and perceived oldness of stimuli from 1 (lowest confidence) to 4 (highest confidence). A prompt with the confidence ratings was displayed for 1.7 seconds. Each trial was separated by an inter-trial interval (ITI) consisting of a fixation cross with a randomly distributed duration of 0-5.4 seconds (mean ITI = 2.7 seconds).

### 2.4 Data Acquisition

MRI scan parameters were matched to those reported in the original publications (Daselaar, Fleck, & Cabeza, 2006; Daselaar, Fleck, Prince, et al., 2006). We collected MRI data using a Siemens Trio 3T MRI scanner (12-channel head coil) and behavioral responses using a four-key fiber-optic response cylinder (Current Designs). We performed structural scanning (256 × 215 matrix, TR 1900 ms, TE 2.26 ms, FOV 250 × 218 mm, 176 slices, 1 mm slice thickness, 0 mm spacing) at the beginning of the scan session and functional scanning (64 × 64 image matrix, TR 1800 ms, TE 31 ms, FOV 240 mm, 34 slices, 3.8 mm slice thickness) during all experimental runs. An MR-compatible LCD monitor displayed stimuli from the head of the bore, which participants viewed through a mirror mounted on the head coil. MRI data are available at https://openneuro.org/datasets/ds004086.

### 2.5 Data Processing

#### 2.5.1 Preprocessing

All fMRI data preprocessing and analyses were performed using the Analysis of Functional Neuroimages (AFNI; Cox, 1996) suite of programs and *fMRIPrep* on BrainLife.io (Avesani et al., 2019). Data were converted from DICOM to NIfTI using dcm2niix, a standalone conversion program (Li et al., 2016). Structural scans were anonymized by removing facial features using in-house scripts prior to being uploaded to OpenNeuro.org for processing with BrainLife.io tools. The description of the preprocessing steps, which included spatial motion correction and spatial normalization, was automatically generated by the *fMRIPrep* software and is included in Appendix 1. Following *fMRIprep* preprocessing, functional data were scaled to have a mean of 100 and blurred with an 8 mm full-width, half-maximum (FWHM) Gaussian kernel to account for inter-subject anatomical variation. Analysis scripts are available here: https://osf.io/ctvsw/.

#### 2.5.2 First Level Regression Analyses

Single-subject regression analyses were conducted using AFNI’s 3dDeconvolve program (Cox, 1996; Cox & Hyde, 1997; Gold et al., 1998). Three regression models were created, all of which included polynomial regressors to account for scanner drift and six motion regressors per run (3 rotations and 3 translations) derived from fMRIPrep output. Task regressors in the first regression analysis coded for the true and perceived oldness of the word and participants’ behavioral responses regardless of confidence (i.e., hits, misses, false alarms, and correct rejections), replicating the analysis of (Daselaar, Fleck, Prince, et al., 2006). In the second regression analysis, regressors coded for behavioral responses and confidence levels independent of the true and perceived oldness of the word. In accordance with (Daselaar, Fleck, & Cabeza, 2006), trials with the lowest confidence level were included in the first-level analysis but then dropped from further analysis resulting in six memory strength levels: 1 = high confidence new to 6 = high confidence old. In the third regression model, only correct memory judgments were included in the model, i.e., all “new” judgments (memory strengths 1, 2, and 3) were correct rejections and all “old” judgments (memory strengths 4, 5, and 6) were hits.

#### 2.5.3 Region of Interest Analysis

In order to determine if the effects from the original papers replicated, we set spherical ROIs (radius of 8 mm) at the peak voxel coordinates of the main activations reported in the previous papers (Guerin & Miller, 2009; O’Connor et al., 2010). Coordinates for spherical ROIs were set in Montreal Neurological Institute (MNI) space as follows based on (Daselaar, Fleck, & Cabeza, 2006): left and right posterior hippocampus (left - 26, -26, -11; right 30, -23, -11), left posterior parahippocampal cortex (−34, -41, -8), left anterior hippocampus (−19, -8, -16), and right rhinal cortex (27, -4, -27). Additionally, based on (Daselaar, Fleck, Prince, et al., 2006), ROIs were placed in bilateral posterior parahippocampal/hippocampal regions (left 27, 27, -15; right -30, 23, -15, respectively). Mean β-coefficients from the single-subject regression analyses were extracted for each ROI using AFNI’s 3dROIstats program and then subjected to repeated-measures ANOVAs as described below.

#### 2.5.4 Exploratory Voxel-Wise Analyses

While the region of interest analyses focused on the coordinates reported in the original studies, heterogeneity in functional localization between subjects may lead to differences in the localization of activations between our sample and the original study sample. Accordingly, we conducted exploratory voxel-wise analyses focusing on voxels only within the MTL. We constructed an MTL mask that included the hippocampus and parahippocampal gyrus bilaterally as defined by segmenting a lab-specific MNI template according to the Desikan-Killiany-Tourville protocol using the Joint Label Fusion toolkit (Klein & Tourville, 2012; Tustison et al., 2014; Wang & Yushkevich, 2013). The high-resolution (1×1×1mm) mask was resampled to the same resolution as the functional data (3.75×3.75×3.8mm) using AFNI program 3dFractionize with a partial-volume clip threshold of 0.8, meaning that 80% or more of the lower-resolution voxel must be occupied to be included in the mask. The subsequent mask was further edited (by author CBK) to exclude extra-MTL voxels (e.g., in lateral ventricles). Voxel-wise analyses were corrected for multiple comparisons first by estimating the smoothness of the functional data from the single-subject regression residuals using AFNI program 3dFWHMx to simulate noise volume where parameters may be estimated (Cox et al., 2017). The smoothness estimates (0.496723, 4.85762, and 10.6138) were incorporated into Monte Carlo simulations with AFNI program 3dClustSim (Cox et al., 2017) using the edited MTL mask volume (1206 voxels in mask), resulting in a spatial extent threshold of k ≥ 12 voxels and voxel-wise p < .001 for a family-wise error corrected p < .01.

We first examined the effects of true and perceived oldness (Daselaar, Fleck, Prince, et al., 2006) using a repeated-measures ANOVA with factors coding for true oldness (targets, foils), perceived oldness (“old”, “new” responses), and sex (male, female). Group-level analyses were performed using AFNI program 3dMVM on the parameter estimates (betas) for Hits, Misses, CRs and FAs from the first-level single-subject analysis. Resulting F maps were thresholded as outlined above. The second analysis examined the effects of recognition confidence using a repeated-measures ANOVA with factors coding for recognition confidence with six levels (Daselaar, Fleck, & Cabeza, 2006) and for sex with 3dMVM. Contrasts were created to code for recollection using a nonlinear function that was flat for levels 1-5 and increased at level 6 (i.e., -1, -1, -1, -1, 5). Familiarity and novelty are graded mnemonic processes and were modeled as contrasts of a continuous linear function. Familiarity was modeled as a positive contrast, in which the level of recognition increases as familiarity increases (i.e., -5, -3, -1, 1, 3, 5). In contrast, novelty was modeled as a negative contrast, in which the level of recognition decreased as novelty-related activations increased (i.e., 5, 3, 1, -1, -3, -5). This analysis identified any voxels in the MTL where activation followed one of the patterns outlined above. Consequently, voxels with activation that roughly increases with increasing recognition confidence may be fit by either function. Accordingly, we followed the analysis in Daselaar, Fleck, and Cabeza (2006) by modeling recollection, familiarity, and novelty as parametric regressors in the first-level single-subject analysis and then directly comparing the resulting fit estimates using paired t-tests at p<0.05 and inclusively masking with the main effect of each contrast thresholded at p<0.1. For example, recollection-related activity was taken as recollection > familiarity masked with the main effect of recollection. We only performed this analysis for recollection and familiarity as the above ANOVA did not reveal any novelty-related signals in the MTL even at liberal thresholds.

## 3. Results

### 3.1 Behavioral Results

The mean number of trials from the test phase across all behavioral responses (i.e., hits, misses, CRs, and FAs) and confidence levels (1 = lowest, 4 = highest) is reported in Table 1. Collapsing across confidence levels, the overall recognition accuracy (d’ score) was 0.84 (SD, 0.43), which was significantly greater than 0 (t(55) = 13.89, p < 0.001, 95% CI [0.696, 0.931], indicating good memory performance. However, performance in our sample was significantly lower than that reported in the original publication (Daselaar, Fleck, & Cabeza, 2006): d’=1.43 (SD=.6); t(65)=4.187, p<.0001, 95%CI [0.31, 8.7].

**Table 1.** Mean Number of Trials (and Standard Deviations) in the Recognition Memory Task

| Behavioral Outcome | Confidence Rating (1-4) |  |  |  |
| --- | --- | --- | --- | --- |
|  | 1 | 2 | 3 | 4 |
| Hit | 5.1 (5.0) | 16.4 (8.7) | 21.9 (8.1) | 34.9 (22.4) |
| Miss | 7.7 (6.3) | 12.7 (6.4) | 12.5 (9.6) | 3.0 (4.1) |
| CR | 9.4 (8.4) | 21.3 (9.3) | 27.3 (12.7) | 12.1 (11.1) |
| FA | 6.6 (6.4) | 16.3 (7.1) | 15.1 (7.9) | 5.5 (7.5) |

In their analysis of memory confidence, the original study authors (Daselaar, Fleck, & Cabeza, 2006) dropped the lowest confidence rating (rating 1 in Table 1) due to low response rates and collapsed targets and foils for the memory strength analysis (levels 1-6). For comparison, we report mean accuracy for memory strength levels 1-6 from our results in Table 2.

**Table 2.**
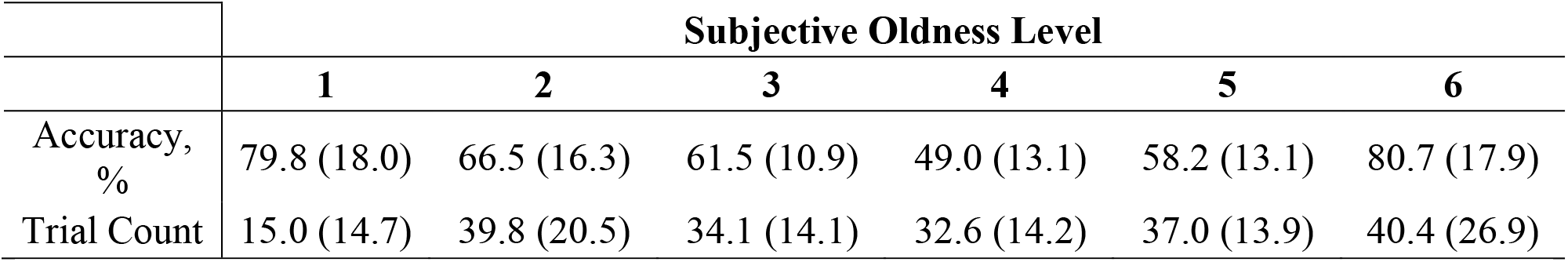
Mean Accuracy (and Standard Deviations) for the Six Subjective Oldness Levels

|  | Subjective Oldness Level |  |  |  |  |  |
| --- | --- | --- | --- | --- | --- | --- |
|  | 1 | 2 | 3 | 4 | 5 | 6 |
| Accuracy, % | 79.8 (18.0) | 66.5 (16.3) | 61.5 (10.9) | 49.0 (13.1) | 58.2 (13.1) | 80.7 (17.9) |
| Trial Count | 15.0 (14.7) | 39.8 (20.5) | 34.1 (14.1) | 32.6 (14.2) | 37.0 (13.9) | 40.4 (26.9) |

### 3.2 ROI Analysis: True vs. Perceived Oldness

Our first set of analyses examined activation for true oldness compared with perceived oldness in ROIs based on (Daselaar, Fleck, Prince, et al., 2006). The original result showed a strong main effect of true oldness with greater activation in posterior MTL regions for both hits and misses compared to CRs and FAs (i.e., targets > foils). We extracted mean β-coefficients from spherical ROIs centered on activation coordinates reported in the original paper (Left: -27, -27, -15; Right: 30, -23, -15; see Figure 2) and subjected them to a repeated-measures ANOVA with factors of hemisphere (left, right), true oldness (targets, foils), and perceived oldness (“old”, “new”). Consistent with the original study, we observed a significant main effect of true oldness collapsed across both hemispheres (F (1, 52) = 10.99, p = 0.002, *η*_*p*_^2^ = 0.174). Consistent with the original work, there was no main effect of perceived oldness (F(1, 52) = 2.31, p = 0.135, *η*_*p*_^2^ = 0.042). There was a significant main effect of hemisphere (F(1,52) = 5.99, p = 0.018, *η*_*p*_^2^ = 0.103) and a significant interaction between hemisphere and perceived oldness (F(1,52) = 12.0, p = .001, *η*_*p*_^2^ = 0.187). Additionally, there was a true oldness by perceived oldness interaction (F(1,52) = 12.678, p < 0.001, *η*_*p*_^2^ = 0.196), see Figure 2. Post-hoc comparisons in the left MTL revealed significant differences between activation for hits and misses (t(52) = -4.394, p < 0.001), hits and CRs (t(52) = -4.727, p < 0.001), and hits and FAs (t(52) = -3.936, p < 0.001). In the right MTL, significant differences were found between hits and misses (t(52) = 3.435, p = 0.001), hits and CRs (t(52) -2.164, p = 0.035), and CRs and misses (t(52) = 2.251, p = 0.029), but not between hits and FA’s. Overall, these results indicate strong activation in the MTL, particularly on the left, for correct “old” decisions (i.e., Hits) compared to other memory outcomes.

**Figure 2.**
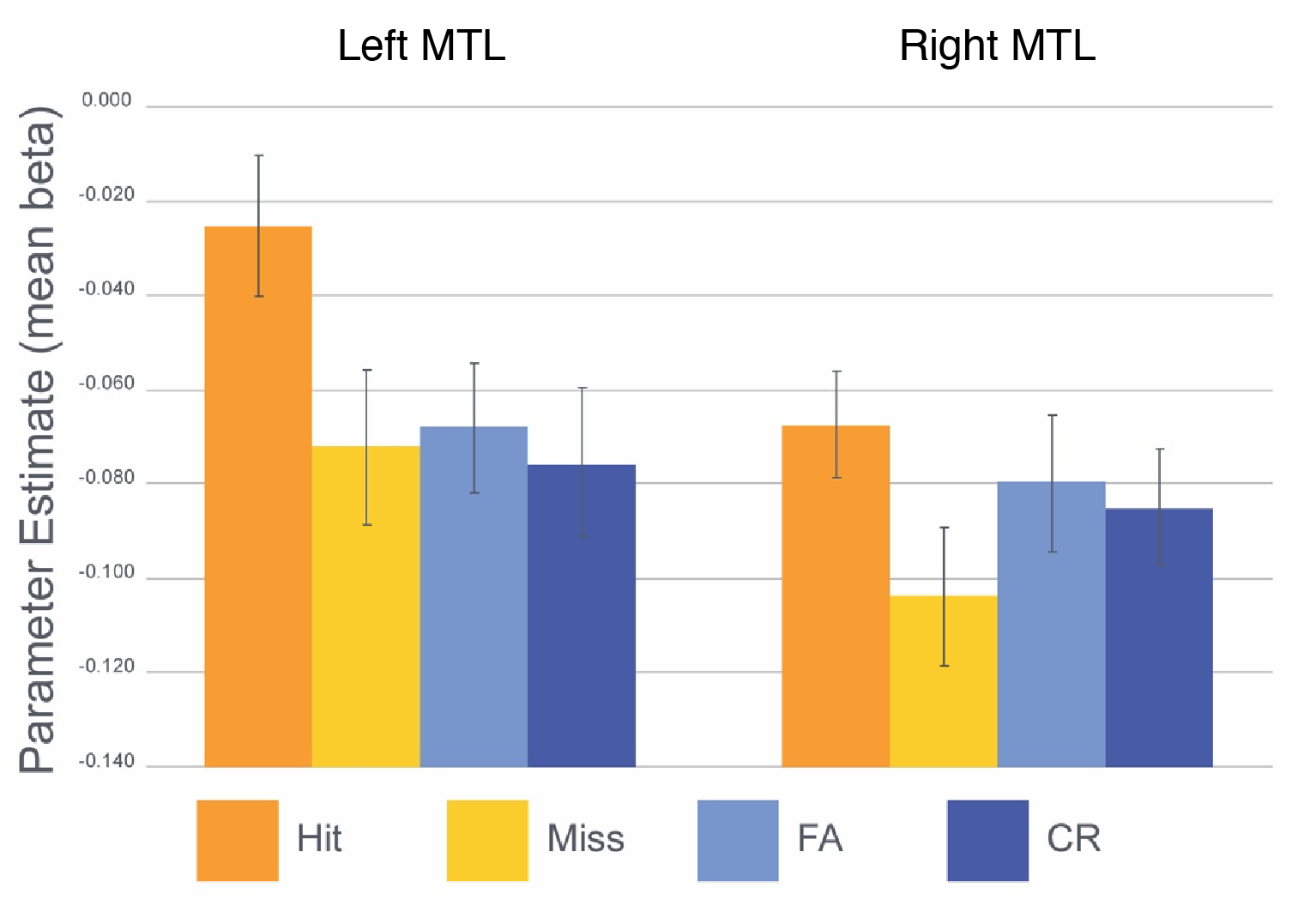
Mean activations related to the discrimination of true and perceived oldness. Error bars indicate +/-SEM. CR = correct rejection; FA = false alarm. The inter-stimulus interval was used as the baseline in the model, resulting in all parameter estimates falling below “baseline” (set as 0 in the model).

### 3.3 ROI Analysis: Subjective Confidence

Our second set of analyses examined activation corresponding to subjective recognition memory confidence based on (Daselaar, Fleck, & Cabeza, 2006). In the previous paper, researchers observed a triple dissociation in which 1) the posterior hippocampus demonstrated a sharp increase in activity for the highest oldness level (consistent with recollection), 2) posterior parahippocampal cortex activity reflected continuous increase as a function of oldness (familiarity), and 3) anterior hippocampal and rhinal regions exhibited a continuous decrease of activity (novelty; Daselaar, Fleck, Prince, et al., 2006). We again examined activation in spherical ROIs defined as the peak coordinates for the largest activation in each of these contrasts. Mean β-coefficients from these ROIs are depicted in Figure 3. In a repeated-measures ANOVA with factors of ROI and memory strength (1-6), we found significant main effects of ROI (F(4,188) = 13.74, p < 0.001, *η*_*p*_^2^ =0.23) and memory strength (F(5,235) = 10.22, p < 0.001, *η*_*p*_^2^= 0.18) and a significant ROI by memory strength interaction (F(20,940) = 4.48, p < 0.001, *η*_*p*_^2^ = 0.09). The significant interaction between brain region and experimental condition in the omnibus ANOVA indicates at least a single dissociation within the MTL. We conducted follow-up analyses to better characterize the activations within the MTL. We found the “recollection” contrast (−1, -1, -1, -1, 5) was significant in each of the *a priori* ROIs (Table 3), with the strongest effect size in left posterior hippocampus (d = 1.14) and the weakest in the right rhinal region (d = 0.32). The “familiarity” contrast (−5, -3, -1, 1, 3, 5) was significant in left and right posterior hippocampus, left parahippocampal cortex, and left anterior hippocampus (p’s < 0.05) but not in the right rhinal region (p = 0.34). The “novelty” contrast was not significant in any *a priori* ROI. Thus, the present study fails to replicate a triple dissociation in the MTL.

**Figure 3.**
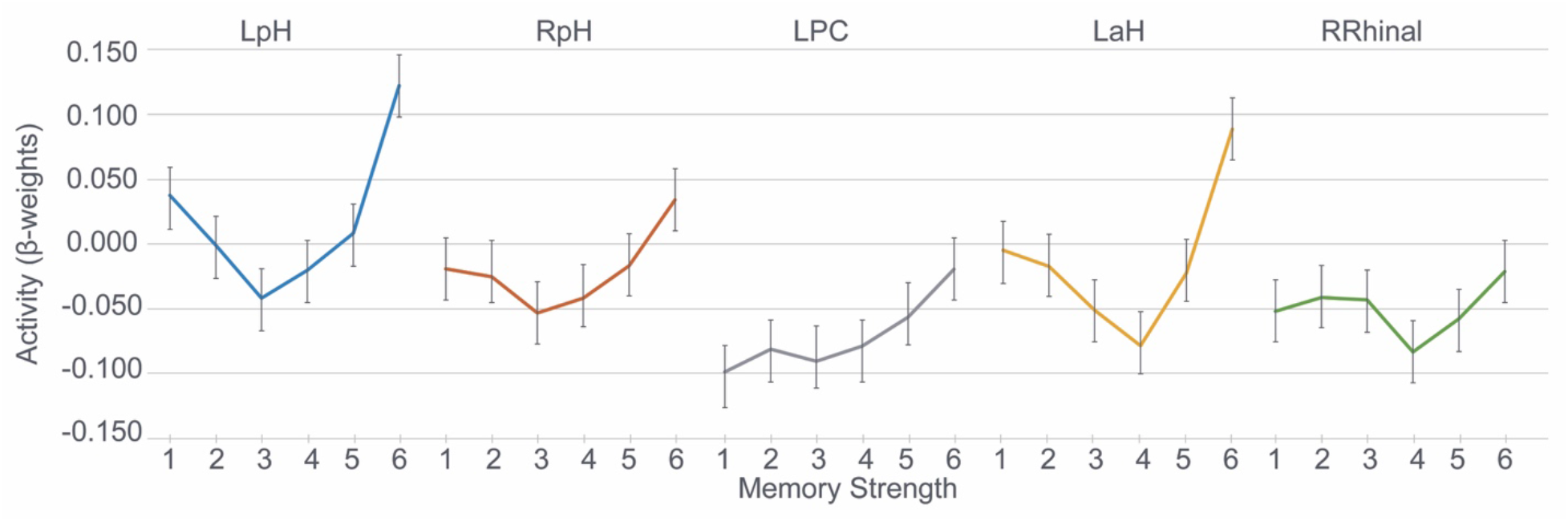
Activations related to Recollection, Familiarity, and Novelty. Error bars indicate +/-SEM. LpH = left posterior hippocampus; RpH = right posterior hippocampus; LPC = Left parahippocampal cortex; LaH = Left anterior hippocampus; RRhinal = right rhinal region.

**Table 3.** Brain Regions Associated with Recollection, Familiarity, and Novelty

| ROI | MNI Coordinates | Effect | t(52) | p (1-tailed) |
| --- | --- | --- | --- | --- |
| LpH | -26, -26, -11 | Familiarity | 4.13 | 6.59E-05 |
|  |  | Novelty | -4.13 | 0.9999 |
|  |  | Recollection | 8.313 | 2.05E-11 |
| RpH | 30, -23, -11 | Familiarity | 3.103 | 0.0015 |
|  |  | Novelty | -3.103 | 0.9985 |
|  |  | Recollection | 4.996 | 3.49E-06 |
| LPHC | -34, -41, -8 | Familiarity | 5.745 | 2.43E-07 |
|  |  | Novelty | -5.745 | 0.9999 |
|  |  | Recollection | 6.274 | 3.55E-08 |
| LaH | -19, -8, -16 | Familiarity | 2.166 | 0.0175 |
|  |  | Novelty | -2.166 | 0.9825 |
|  |  | Recollection | 5.159 | 1.97E-06 |
| RRhinal | 27, -4, -27 | Familiarity | 0.419 | 0.3385 |
|  |  | Novelty | -0.419 | 0.6615 |
|  |  | Recollection | 2.341 | 0.0115 |

*Note:* LpH = left posterior hippocampus; RpH = right posterior hippocampus; LPC = Left parahippocampal cortex; LaH = Left anterior hippocampus; RRhinal = right rhinal region.

### 3.4 SPM Analysis

In order to rule out analysis software choice as an explanation for failure to replicate specific findings of the previous papers, we repeated the above analysis using SPM12. We followed the default preprocessing pipeline in SPM for slice time correction, motion correction, smoothing (8mm FWHM Gaussian), and spatial normalization to the MNI template. We conducted two first-level GLM analyses: one for true vs. perceived oldness analysis (with regressors for hits, misses, CRs, and FAs) and one for the subjective confidence analysis. Significant effects were exactly as in the AFNI analysis above with the exception that the main effect of true oldness in the left MTL failed to reach statistical significance with the SPM pipeline. Since the SPM and AFNI results were materially similar (and indeed the AFNI results replicated slightly better), we opted to use AFNI for the remaining analyses.

### 3.5 Exploratory Voxel-Wise Analyses

To characterize further the overall response to recognition memory tasks in the MTL, we performed two sets of repeated-measures ANOVAs to identify regions with significant differences in activation between true oldness (targets, foils) and perceived oldness (“old”, “new” responses) conditions and then between variations of memory strength (1-6). Sex was included as a factor in both sets of analyses but no main effects or interactions with this factor were observed.

In the true vs. perceived oldness analysis, we observed a cluster in left posterior MTL (Table 4) with greater activation for “old” responses than for “new” responses (mean old = -0.094, mean new = -0.131; t(52) = -5.46, p < 0.001, 95%CI [-0.053, -0.024]). We also observed clusters in left and right posterior parahippocampal cortex where there was an objective by subjective old/new interaction (Figure 4A; Table 4). Consistent with the ROI analysis above, there was more activation for hits than any other condition in each cluster (p’s <0.001).

**Table 4.** Clusters of Activation in MTL Voxel-Wise Analyses

| Contrast | Label | Cluster Size<br>(voxels) | Peak Activation<br>(MNI) |  |  |
| --- | --- | --- | --- | --- | --- |
|  |  |  | X | Y | Z |
| Main Effect of Subjective Response |  |  |  |  |  |
|  | L. Posterior MTL | 42 | -29 | -43 | -6 |
| Subjective x Objective Interaction |  |  |  |  |  |
|  | L. Posterior PHC | 30 | -22 | -43 | -18 |
|  | R. Posterior PHC | 13 | 16 | 31 | -10 |
| Main Effect of Familiarity |  |  |  |  |  |
|  | L. Posterior PHC | 129 | -7 | -39 | 1 |
|  | R. Posterior PHC | 70 | 24 | -43 | -18 |
| Main Effect of Recollection |  |  |  |  |  |
|  | L. MTL | 187 | 9 | -39 | 1 |
|  | R. MTL | 29 | 24 | -39 | -18 |
| Familiarity > Recollection |  |  |  |  |  |
|  | R. Posterior PHC | 32 | -29 | 10 | -41 |
| Recollection > Familiarity |  |  |  |  |  |
|  | L. Hippocampus & ERC | 13 | 16 | 31 | -10 |

*Note:* L. = left; R. = right; PHC = parahippocampal cortex; MTL = medial temporal lobe; ERC = entorhinal cortex

**Figure 4.**
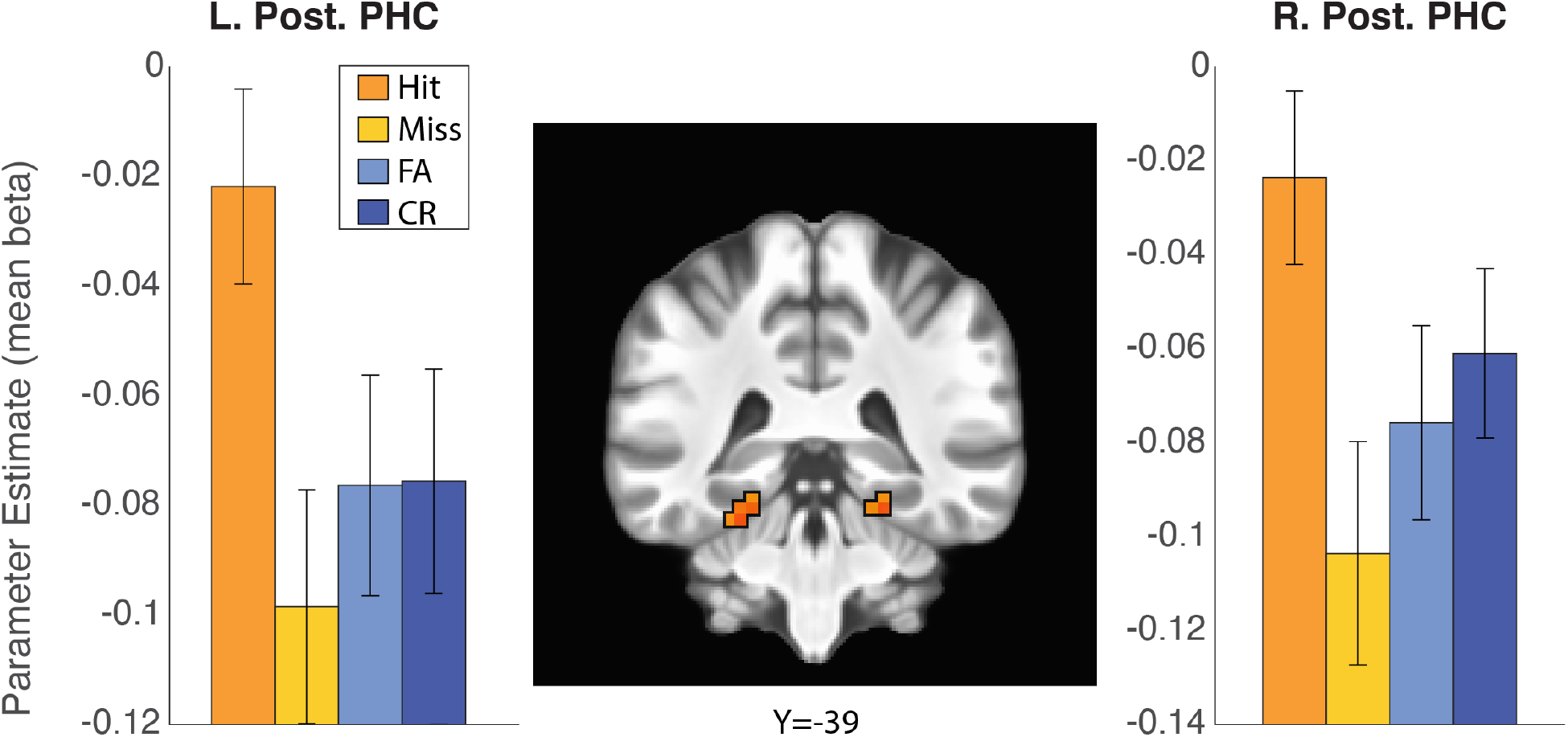
Clusters showing a significant subjective by objective oldness interaction were located in left and right posterior parahippocampal cortex. In each cluster, activation for hits was significantly greater than activation for every other condition. Bar graphs depict mean activation for all voxels within the corresponding cluster. Error bars indicate +/-SEM; L = left; R = right; Post. = posterior; PHC = parahippocampal cortex; Coronal slice location is in MNI coordinates.

In the subjective confidence analysis, we observed large clusters of activation for the familiarity contrast in the parahippocampal gyrus bilaterally (Figure 5A-C; Table 4). We also observed large clusters for the recollection contrast that overlapped substantially with the familiarity clusters but also included more voxels in the hippocampus (Figure 5D-F). We did not observe any voxels that demonstrated a novelty effect, even at a relaxed voxel-wise p<.05.

**Figure 5.**
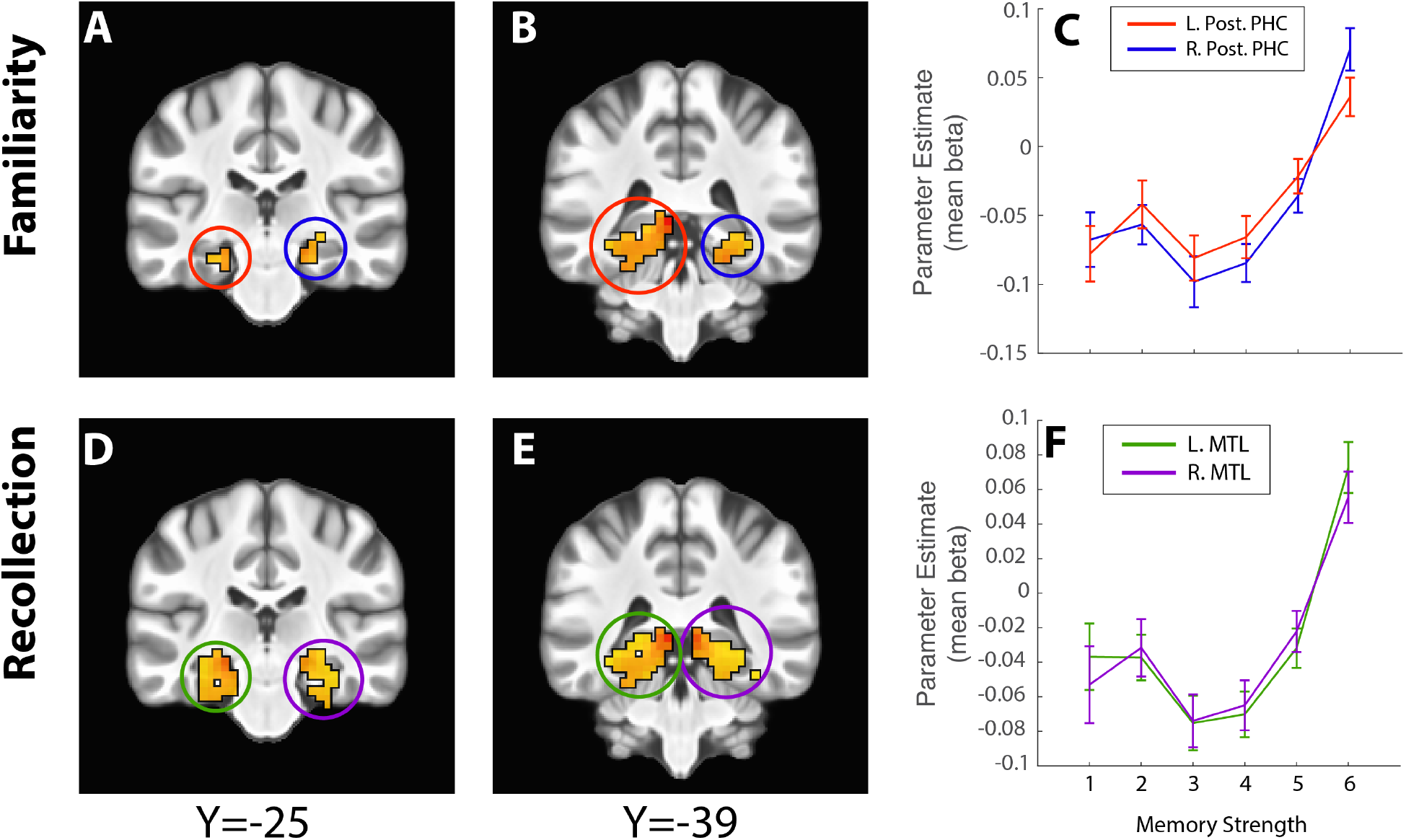
Voxel-wise analysis of familiarity and recollection contrasts. There were large clusters in the left and right posterior parahippocampal cortex (A, B) where the familiarity contrast was significant (C). Similarly, there were large clusters in left and right MTL (D, E) where the recollection contrast was significant (F) that overlapped with the familiarity clusters. Line graphs depict mean activation across all voxels in the cluster and colors in C and F correspond to the highlighting circles in A,B and D,E, respectively. Error bars indicate +/-SEM; L = left; R = right; Post. = posterior; PHC = parahippocampal cortex; lat. = lateral; med. = medial; Coronal slice locations in (A,B,D & E) are in MNI coordinates.

For completeness, we also performed the above voxel-wise analyses for the whole brain data. Since the whole-brain analysis is beyond the scope of this replication attempt, we have posted the whole-brain contrast maps at https://neurovault.org/collections/12889/.

The MTL voxel-wise analysis identified large overlapping clusters in the MTL that were fit by both the recollection and the familiarity contrasts. To give ourselves the best chance at identifying voxels that differentially responded to recollection and familiarity, we followed the analysis of Daselaar, Fleck, and Cabeza (2006) by first identifying voxels that were significantly better fit by either the recollection or familiarity contrast and inclusively masking those clusters with the relevant main effect. This analysis revealed a cluster in posterior right parahippocampal cortex where familiarity > recollection (Figure 6A-B) and a cluster in left anterior MTL that included voxels in the hippocampal head but also anterior entorhinal cortex where recollection > familiarity (Figure 6C-D). The clusters from the familiarity contrast correspond well with those identified in the original paper (Table 4); however, the clusters identified in the recollection analysis are more anterior in our sample and include voxels outside the hippocampus including in the entorhinal cortex (e.g., Figure 6C).

**Figure 6.**
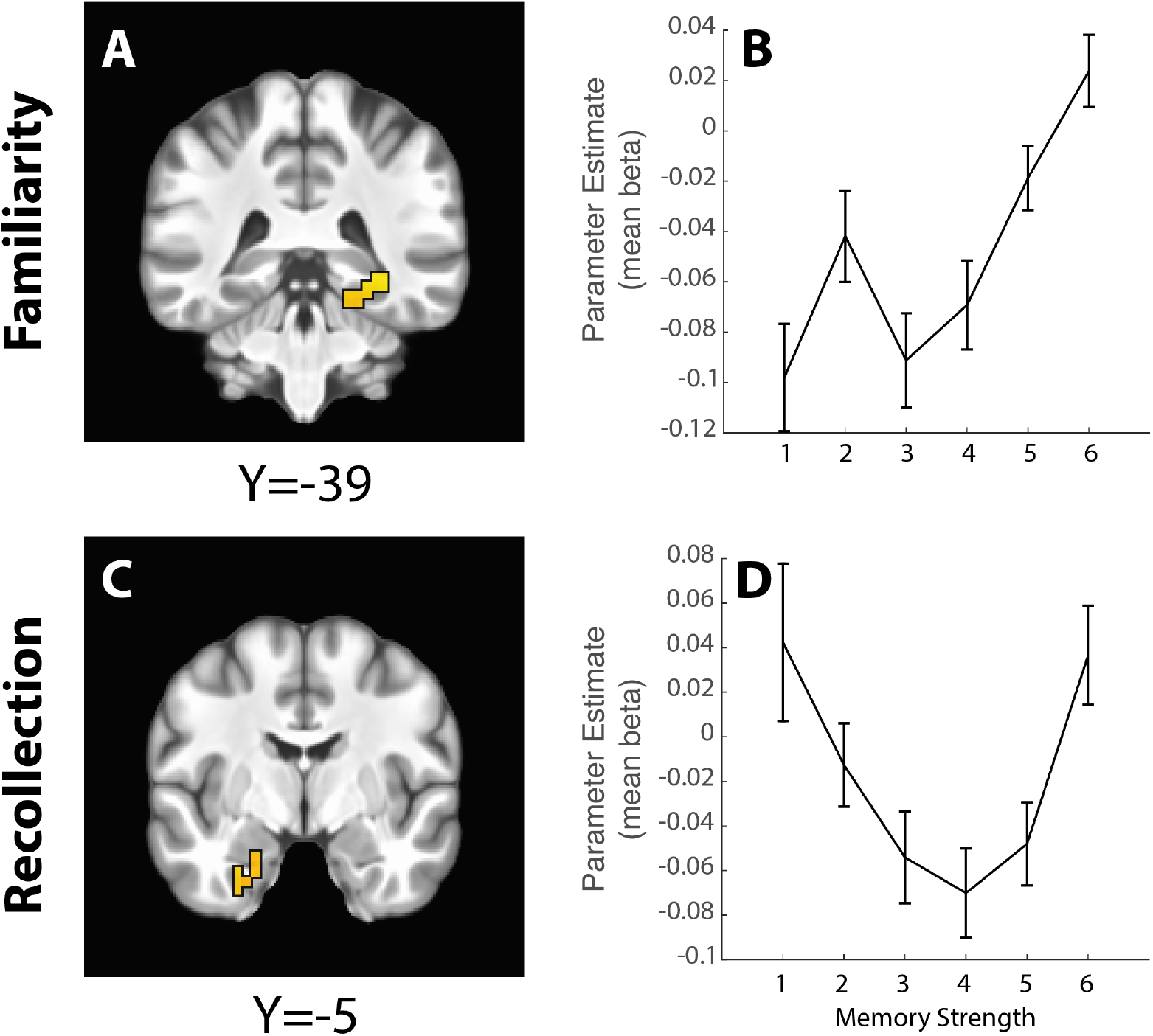
Voxel-wise analysis of Familiarity > Recollection (A, B) and Recollection > Familiarity (C, D) contrasts. Line graphs depict mean activation across all voxels in the corresponding cluster. Error bars indicate +/-SEM; L = left; R = right; Post. = posterior; PHC = parahippocampal cortex; Hip = hippocampus; ERC = entorhinal cortex; Coronal slice locations in (A,B,D & E) are in MNI coordinates.

## 4. Discussion

Here we present an attempt to replicate two papers that performed complementary analyses on a recognition memory fMRI dataset. Our replication efforts were met with mixed success. When considering the MTL response to veridical or true oldness we found the greatest response to hits, with this activation driving all main effects and interactions in *a priori* MTL ROIs defined based on the previously reported activations. This is in contrast to the original finding, in which MTL activation was increased for both hits and misses (true old stimuli) relative to all new stimuli (Daselaar, Fleck, & Cabeza, 2006). The ROI analysis may have been insensitive to effects happening elsewhere in the MTL. Accordingly, we performed a voxel-wise analysis within the MTL and again identified clusters of activation in posterior parahippocampal cortex and retrosplenial cortex where activation for hits was greater than for any other recognition memory outcome. The results of the original paper were taken as evidence that the MTL distinguishes between old and new stimuli regardless of the participants’ subjective experience. In contrast, in our sample the MTL response depended on both the true oldness of the stimulus and the participants’ reported conscious experience and was driven by correct “old” decisions, i.e., hits.

When considering the MTL response to recognition confidence (regardless of true and perceived oldness of the stimulus), we again failed to exactly replicate the results of the previous study (Daselaar, Fleck, & Cabeza, 2006). When considering activation within ROIs based on the reported coordinates of activations in the previous paper, we found strong “recollection” signals in bilateral posterior and left anterior hippocampus, with sharply increased activation at the highest recognition confidence level similar to the original finding. We found “familiarity” signals in the left parahippocampal cortex, with a more linear increase in activation with recognition confidence. These results are consistent with those of the previous study. However, in contrast to the original study, there was no evidence of a “novelty” signal in the rhinal region that decreased with recognition confidence. Again, we performed a voxel-wise analysis to identify recollection, familiarity, and novelty signals in the broader MTL beyond the *a priori* regions of interest. In this broader analysis, we found large clusters in bilateral MTL, mostly contained within parahippocampal cortex (Figure 5A-B), where activation increased linearly as a function of recognition memory strength. We also observed large clusters in bilateral MTL where the recollection contrast was significant. The recollection clusters overlapped substantially with the familiarity clusters (Figure 5D-E) but included more voxels in the hippocampus. Given this high degree of overlap it is difficult to conclude that there is a dissociation between recollection- and familiarity-related responses within the MTL.

This overlap between “recollection” and “familiarity” clusters is, at least in part, driven by the fact that the contrasts used to define recollection and familiarity are not orthogonal to each other. In order to identify regions that differentially responded to recollection and familiarity, we followed the analysis path of Daselaar, Fleck, and Cabeza (2006) by directly contrasting fit coefficients for the two contrasts and then masking the resulting maps with the relevant main effects. In the case of the familiarity > recollection contrast, we observed a cluster of activation in right posterior parahippocampal cortex (Figure 6A) that is similarly located to that reported in the original paper (Daselaar, Fleck, and Cabeza, 2006; Figure 1). In the case of the recollection > familiarity contrast, we identify clusters in anterior MTL that include voxels within the hippocampus (consistent with the original paper) but also include voxels in entorhinal cortex. The previous paper only reported clusters in posterior hippocampus that demonstrated a recollection effect, which we fail to replicate in our voxel-wise analysis.

Taking together all the results of the subjective memory confidence analyses, we at best observe a single dissociation (Henson, 2006) between recollection and familiarity signals within the posterior MTL, similar to other studies (Ranganath et al., 2004). The strongest replication in the present study was the familiarity > recollection signal in the posterior parahippocampal cortex. Given that we did not observe any activation in the MTL consistent with novelty signals even at liberal thresholds, we conclusively fail to replicate a triple dissociation in the MTL.

Several models have been put forward to explain how the MTL supports recognition memory performance. The dual-process signal detection (DPSD) model posits that recollection is critically dependent on the hippocampus due to its role in forming and retrieving associations while familiarity depends on MTL cortex (Yonelinas, 1994). In contrast, a multi-attribute perspective views recollection as more of a continuum of memory confidence that is not merely limited to the highest level of confidence (Wixted & Squire, 2011). This model, which proposes that the hippocampus contributes to both recollection and familiarity processes, has received mixed support in how recognition memory tests have been interpreted and in how the hippocampus contributes to these processes (Carr et al., 2010; Jenkins et al., 2004). The multi-attribute theory suggests that while high-confidence responses are considered recollection, low-confidence responses reflect familiarity in a graded effect across a continuum. A more recent alternative model attempts to distinguish between use of single-versus multi-attribute stimuli, suggesting that outcomes may differ depending on the attributional features of stimuli in experimentation (i.e., a stimulus with a single attribute may link recollection and familiarity to hippocampal functioning while a stimulus with multiple attributes may reveal that familiarity is instead supported by other MTL structures; Marzolla, 2018). Findings from the present study are consistent with this recent proposal in that we observed hippocampal activations associated with both recollection and familiarity (Table 3). According to this model, this is likely due to the single attributional features of the stimuli used; in this case, words. Implications from the alternative model argue for a shift in methodology when studying recognition memory in future work by, for example, involving the use of multidimensional stimuli such as neutral faces as single attributional stimuli and emotional faces for multiple attributional stimuli.

Prior research has shown stimulus-specific responding across regions within the MTL. Specifically, Kafkas and colleagues (Kafkas et al., 2017) found material specificity drove familiarity-based responses within the MTL but not recollection within the hippocampus. Select MTL regions (perirhinal and entorhinal cortices) were responsive to object familiarity while others (parahippocampal cortex) responded to both object and scene familiarity. In contrast, the hippocampus was involved in recollection independent of the stimulus type (Kafkas et al., 2017). Both these findings and distinctions made from the alternative model of hippocampal and MTL functioning in recognition memory suggest that the specificity of the stimuli may also be influencing how each region is involved.

Other studies have found recollection-based representations in the hippocampus to be more flexible than familiarity-based representations in MTL structures like the perirhinal and parahippocampal cortices (Ozubko et al., 2017). These differences are driven by how recollection- and familiarity-based associations are formed and supported, as recollection-based representations are discretely coded and bound together for later learning (through the hippocampus) and familiarity-based representations are more susceptible to interference given the challenge of distinguishing elements from past associations (through the MTL; Ozubko et al., 2017). Given the high variability of support among competing models of recognition memory to date, we again emphasize the importance of replication to further clarify the roles between the hippocampus and MTL structures in recollection and familiarity processes, lending more support on some models than for others.

One potential concern when organizing trials by subjective recognition confidence responses without consideration for accuracy is that the highest confidence category tends to have mostly correct trials while the proportion of correct trials decreases in lower confidence categories. Previous studies have noted that this may result in selective averaging that could account for “recollection” activation patterns like the one we observed in the hippocampus (Wixted & Squire, 2011). To rule out this account we repeated our analyses of the memory confidence data using only correct responses (i.e., hits and correct rejections separated by high, medium, and low confidence) and dropping incorrect trials. The pattern of results with only correct trials was the same as when we analyzed the full data set. Thus, the results were remarkably similar and indicating no influence of selective averaging.

Another consideration when collapsing across veridical old/new status is potential encoding-related activation potentially interfering with retrieval-related activation. One would assume that encoding related activation would be strongest for the lowest memory strength bins (i.e., 1-3) but lower for higher bins. Given the strong pattern of increasing activation with increasing memory strength levels, we do not think this is a major confound in our data, but it cannot be ruled out entirely.

While we attempted to follow the original research as exactly as possible there were a few unavoidable differences between the original experiments and the present replication study. To match the study population, the present study recruited from a university community much like the population used in the previous papers, which did not report additional demographics beyond age and sex. Ages of the two samples were approximately matched (21.7 vs. 21.3). In the original study, 6 of 14 (43%) of participants were women, while 33 of 53 (62%) were women in our sample. We included sex as a factor in all analyses but did not observe any main effects or interactions with sex in any of our results, indicating that any difference in the distribution of males and females across the samples did not affect the results. While we endeavored to match study materials as closely as possible, differences in materials may account for performance differences between the studies and potentially may account for activation differences as well. Further, MRI equipment and scan parameters differed from the original study. Although we again attempted to match the parameters reported originally, some differences were unavoidable. Scan sequences and preprocessing routines were made available as open source for future replications (e.g., posted to brainlife.io). Additionally, we used an ROI-based approach rather than a voxel-based approach as in the original studies. This approach was adopted to avoid potential differences introduced by different procedures for voxel-based analyses across software packages (e.g., SPM2 vs. AFNI) in order to perform rigorous tests for replication. Other differences, however, were largely due to efforts to improve the ease of replication and study methodology. Stimuli, data, and scripts from the present study have been made available for download and access so that other labs may replicate our analyses. The fMRIPrep boilerplate used in preprocessing our data is available in the Appendix. While Daselaar et al. used an SPM preprocessing pipeline, repeating our analyses with SPM did not produce appreciably different results. Indeed, we hope that using the latest and most advanced/open tools to perform our analyses may provide clearer data, resulting in less noise and more signal.

Our sample size was determined by an *a priori* power analysis based on the estimated effect size of the “true oldness” effect in the right posterior MTL (Daselaar, Fleck, Prince, et al., 2006; Figure 1B). This effect size was chosen as it appeared to be the smallest effect size reported across both original papers. The intent was to avoid overly optimistic power calculations (Anderson & Maxwell, 2017), but our sample size may well have still been underpowered to detect small effects (Szucs & Ioannidis, 2020).

In conclusion, our well-powered study found a consistent role of the parahippocampal region in the subjective experience of remembering in a recognition memory paradigm. When only considering whether yes/no recognition memory decisions were correct, only accurate “old” decisions (i.e., hits) were associated with increased parahippocampal responses. When considering the subjective experience of recognition confidence, we found increased parahippocampal activation for the highest confidence level and a more graded response in the MTL cortex, consistent with a role of these areas in recollection and a role of the MTL cortex in familiarity.

While past works may be strongly influential from a theoretical perspective as to what the hippocampus does in relation to memory, neuroimaging research has often involved features thought to yield high levels of false findings. A high degree of flexibility in neuroimaging analyses also leave studies vulnerable to the inflation of false-positive rates. Here, while we demonstrate that essential features of the original publications do replicate, we were not able to replicate all findings (e.g., no novelty signal in the MTL; no evidence of increased parahippocampal activation for old stimuli identified as “new” or misses). The present work highlights the necessity of replication as a routine procedure that must be integrated into science, especially in neuroimaging research, as a means to critically review and confirm outcomes as they are published and used for other works.

## 5. Acknowledgements

We thank Alice Wang, Isaac Hamm, Austin Schmidt, Sydney Willden, Morgan Chase, and Clara Rodriguez for their support on study design, data collection, and quality assurance. We thank Arianna Hedges-Muncy for her assistance with *a priori* power analyses. We also extend our acknowledgements to Dr. Steven Allen for his technical support and consultation. This work was supported by the College of Family, Home, and Social Sciences Mentored Environment Grant (MEG) at Brigham Young University.

## Appendix A: fMRIprep Description

Results included in this manuscript come from preprocessing performed using *fMRIPrep* 20.2.1 (@fmriprep1; @fmriprep2; RRID:SCR_016216), which is based on *Nipype* 1.5.1 (@nipype1; @nipype2; RRID:SCR_002502).

### Anatomical Data Preprocessing

The T1-weighted (T1w) image was corrected for intensity non-uniformity (INU) with N4BiasFieldCorrection (@n4), distributed with ANTs 2.3.3 (@ants, RRID:SCR_004757), and used as T1w-reference throughout the workflow. The T1w-reference was then skull-stripped with a *Nipype* implementation of the antsBrainExtraction.sh workflow (from ANTs), using NKI as target template. Brain tissue segmentation of cerebrospinal fluid (CSF), white-matter (WM) and gray-matter (GM) was performed on the brain-extracted T1w using fast (FSL 5.0.9, RRID:SCR_002823, @fsl_fast). Brain surfaces were reconstructed using recon-all (FreeSurfer 6.0.1, RRID:SCR_001847, @fs_reconall), and the brain mask estimated previously was refined with a custom variation of the method to reconcile ANTs-derived and FreeSurfer-derived segmentations of the cortical gray-matter of Mindboggle (RRID:SCR_002438, @mindboggle). Volume-based spatial normalization to two standard spaces (MNI152NLin2009cAsym, MNI152NLin6Asym) was performed through nonlinear registration with antsRegistration (ANTs 2.3.3), using brain-extracted versions of both T1w reference and the T1w template. The following templates were selected for spatial normalization: *ICBM 152 Nonlinear Asymmetrical template version 2009c* (@mni152nlin2009casym, RRID:SCR_008796; TemplateFlow ID: MNI152NLin2009cAsym), *FSL’s MNI ICBM 152 non-linear 6th Generation Asymmetric Average Brain Stereotaxic Registration Model* (@mni152nlin6asym, RRID:SCR_002823; TemplateFlow ID: MNI152NLin6Asym).

### Functional Data Preprocessing

For each of the BOLD runs found for each participant (across all tasks and sessions), the following preprocessing was performed. First, a reference volume and its skull-stripped version were generated using a custom methodology of *fMRIPrep*. A deformation field to correct for susceptibility distortions was estimated based on *fMRIPrep*’s *fieldmap-less* approach. The deformation field is that resulting from co-registering the BOLD reference to the same-subject T1w-reference with its intensity inverted (@fieldmapless1; @fieldmapless2). Registration is performed with antsRegistration (ANTs 2.3.3), and the process regularized by constraining deformation to be nonzero only along the phase-encoding direction and modulated with an average fieldmap template (@fieldmapless3). Based on the estimated susceptibility distortion, a corrected EPI (echo-planar imaging) reference was calculated for a more accurate co-registration with the anatomical reference. The BOLD reference was then co-registered to the T1w reference using bbregister (FreeSurfer) which implements boundary-based registration (@bbr). Co-registration was configured with six degrees of freedom. Head-motion parameters with respect to the BOLD reference (transformation matrices, and six corresponding rotation and translation parameters) are estimated before any spatiotemporal filtering using mcflirt (FSL 5.0.9, @mcflirt). BOLD runs were slice-time corrected using 3dTshift from AFNI 20160207 (@afni, RRID:SCR_005927). The BOLD time-series were resampled onto the following surfaces (FreeSurfer reconstruction nomenclature): *fsaverage*. The BOLD time-series (including slice-timing correction when applied) were resampled onto their original, native space by applying a single, composite transform to correct for head-motion and susceptibility distortions. These resampled BOLD time-series will be referred to as *preprocessed BOLD in original space*, or just *preprocessed BOLD*. The BOLD time-series were resampled into standard space, generating a *preprocessed BOLD run in MNI152NLin2009cAsym space*. First, a reference volume and its skull-stripped version were generated using a custom methodology of *fMRIPrep. Grayordinates* files (@hcppipelines) containing 91k samples were also generated using the highest-resolution fsaverage as intermediate standardized surface space. Several confounding time-series were calculated based on the *preprocessed BOLD*: framewise displacement (FD), DVARS and three region-wise global signals. FD was computed using two formulations following Power (absolute sum of relative motions, @power_fd_dvars) and Jenkinson (relative root mean square displacement between affines, @mcflirt). FD and DVARS are calculated for each functional run, both using their implementations in *Nipype* (following the definitions by @power_fd_dvars). The three global signals are extracted within the CSF, the WM, and the whole-brain masks. Additionally, a set of physiological regressors were extracted to allow for component-based noise correction (*CompCor*, @compcor). Principal components are estimated after high-pass filtering the *preprocessed BOLD* time-series (using a discrete cosine filter with 128s cut-off) for the two *CompCor* variants: temporal (tCompCor) and anatomical (aCompCor). tCompCor components are then calculated from the top 2% variable voxels within the brain mask. For aCompCor, three probabilistic masks (CSF, WM and combined CSF+WM) are generated in anatomical space. The implementation differs from that of Behzadi et al. in that instead of eroding the masks by 2 pixels on BOLD space, the aCompCor masks are subtracted a mask of pixels that likely contain a volume fraction of GM. This mask is obtained by dilating a GM mask extracted from the FreeSurfer’s *aseg* segmentation, and it ensures components are not extracted from voxels containing a minimal fraction of GM. Finally, these masks are resampled into BOLD space and binarized by thresholding at 0.99 (as in the original implementation). Components are also calculated separately within the WM and CSF masks. For each CompCor decomposition, the *k* components with the largest singular values are retained, such that the retained components’ time series are sufficient to explain 50 percent of variance across the nuisance mask (CSF, WM, combined, or temporal). The remaining components are dropped from consideration. The head-motion estimates calculated in the correction step were also placed within the corresponding confounds file. The confound time series derived from head motion estimates and global signals were expanded with the inclusion of temporal derivatives and quadratic terms for each (@confounds_satterthwaite_2013). Frames that exceeded a threshold of 0.5 mm FD or 1.5 standardised DVARS were annotated as motion outliers. All resamplings can be performed with *a single interpolation step* by composing all the pertinent transformations (i.e. head-motion transform matrices, susceptibility distortion correction when available, and co-registrations to anatomical and output spaces). Gridded (volumetric) resamplings were performed using antsApplyTransforms (ANTs), configured with Lanczos interpolation to minimize the smoothing effects of other kernels (@lanczos). Non-gridded (surface) resamplings were performed using mri_vol2surf (FreeSurfer).

Many internal operations of *fMRIPrep* use *Nilearn* 0.6.2 (@nilearn, RRID:SCR_001362), mostly within the functional processing workflow. For more details of the pipeline, see <u>the section corresponding to workflows in</u> *<u>fMRIPrep</u>*<u>’s documentation</u>.

## Copyright Waiver

The above boilerplate text was automatically generated by fMRIPrep with the express intention that users should copy and paste this text into their manuscripts *unchanged*. It is released under the <u>CC0</u> license.

## Author’s Contributions

All authors contributed equally to this work. J.V. prepared the manuscript, implemented study procedures, performed data analyses, and interpreted replication outcomes (under the guidance of the second author). C.B.K. conceived the replication and designed the methodology with the support of his lab (listed in Acknowledgements). All authors read and approved the final manuscript.

## References

Anderson, S. F., & Maxwell, S. E. (2017). Addressing the “Replication Crisis”: Using Original Studies to Design Replication Studies with Appropriate Statistical Power. Multivariate Behavioral Research, 52(3), 305–324. https://doi.org/10.1080/00273171.2017.1289361

Avesani, P., McPherson, B., Hayashi, S., Caiafa, C. F., Henschel, R., Garyfallidis, E., Kitchell, L., Bullock, D., Patterson, A., Olivetti, E., Sporns, O., Saykin, A. J., Wang, L., Dinov, I., Hancock, D., Caron, B., Qian, Y., & Pestilli, F. (2019). The open diffusion data derivatives, brain data upcycling via integrated publishing of derivatives and reproducible open cloud services. Scientific Data, 6(1), 69. https://doi.org/10.1038/s41597-019-0073-y

Baker, M. (2016). 1,500 scientists lift the lid on reproducibility. Nature, 533(7604), 452–454. https://doi.org/10.1038/533452a

Button, K. S., Ioannidis, J. P., Mokrysz, C., Nosek, B. A., Flint, J., Robinson, E. S., & Munafo, M. R. (2013). Power failure: Why small sample size undermines the reliability of neuroscience. Nat Rev Neurosci, 14(5), 365–376. https://doi.org/10.1038/nrn3475

Carr, V. A., Viskontas, I. V., Engel, S. A., & Knowlton, B. J. (2010). Neural Activity in the Hippocampus and Perirhinal Cortex during Encoding Is Associated with the Durability of Episodic Memory. Journal of Cognitive Neuroscience, 22(11), 2652–2662. https://doi.org/10.1162/jocn.2009.21381

Cox, R. W. (1996). AFNI: Software for analysis and visualization of functional magnetic resonance neuroimages. Computers and Biomedical Research, an International Journal, 29(3), 162–173. https://doi.org/10.1006/cbmr.1996.0014

Cox, R. W., Chen, G., Glen, D. R., Reynolds, R. C., & Taylor, P. A. (2017). FMRI Clustering in AFNI: False-Positive Rates Redux. Brain Connect, 7(3), 152–171. https://doi.org/10.1089/brain.2016.0475

Cox, R. W., & Hyde, J. S. (1997). Software tools for analysis and visualization of fMRI data. NMR in Biomedicine, 10(4–5), 171–178. https://doi.org/10.1002/(sici)1099-1492(199706/08)10:4/5<171::aid-nbm453>3.0.co;2-l

Current Designs. (n.d.). 4 Side Button Stick (Cylinder) Fiber Optic Response Pad (900 series) [Computer software].

Daselaar, S. M., Fleck, M. S., & Cabeza, R. (2006). Triple dissociation in the medial temporal lobes: Recollection, familiarity, and novelty. J Neurophysiol, 96(4), 1902–1911. https://doi.org/10.1152/jn.01029.2005

Daselaar, S. M., Fleck, M. S., Prince, S. E., & Cabeza, R. (2006). The medial temporal lobe distinguishes old from new independently of consciousness. J Neurosci, 26(21), 5835–5839. https://doi.org/26/21/5835 [pii] 10.1523/JNEUROSCI.0258-06.2006

Faul, F., Erdfelder, E., Buchner, A., & Lang, A.-G. (2009). Statistical power analyses using G*Power 3.1: Tests for correlation and regression analyses. Behavior Research Methods, 41(4), 1149–1160. https://doi.org/10.3758/BRM.41.4.1149

Gold, S., Christian, B., Arndt, S., Zeien, G., Cizadlo, T., Johnson, D. L., Flaum, M., & Andreasen, N. C. (1998). Functional MRI statistical software packages: A comparative analysis. Human Brain Mapping, 6(2), 73–84.

Guerin, S. A., & Miller, M. B. (2009). Lateralization of the parietal old/new effect: An event-related fMRI study comparing recognition memory for words and faces. NeuroImage, 44(1), 232–242. https://doi.org/10.1016/j.neuroimage.2008.08.035

Henson, R. (2006). Forward inference using functional neuroimaging: Dissociations versus associations. Trends in Cognitive Sciences, 10(2), 64–69. https://doi.org/10.1016/j.tics.2005.12.005

Jenkins, T. A., Amin, E., Pearce, J. M., Brown, M. W., & Aggleton, J. P. (2004). Novel spatial arrangements of familiar visual stimuli promote activity in the rat hippocampal formation but not the parahippocampal cortices: A c-fos expression study. Neuroscience, 124(1), 43–52. https://doi.org/10.1016/j.neuroscience.2003.11.024

Kafkas, A., Migo, E. M., Morris, R. G., Kopelman, M. D., Montaldi, D., & Mayes, A. R. (2017). Material Specificity Drives Medial Temporal Lobe Familiarity But Not Hippocampal Recollection. Hippocampus, 27(2), 194–209. https://doi.org/10.1002/hipo.22683

Kirwan, C. B., Shrager, Y., & Squire, L. R. (2009). Medial temporal lobe activity can distinguish between old and new stimuli independently of overt behavioral choice. Proceedings of the National Academy of Sciences, 106(34), 14617–14621. https://doi.org/10.1073/pnas.0907624106

Klein, A., & Tourville, J. (2012). 101 labeled brain images and a consistent human cortical labeling protocol. Front Neurosci, 6, 171. https://doi.org/10.3389/fnins.2012.00171

Li, X., Morgan, P. S., Ashburner, J., Smith, J., & Rorden, C. (2016). The first step for neuroimaging data analysis: DICOM to NIfTI conversion. Journal of Neuroscience Methods, 264, 47–56. https://doi.org/10.1016/j.jneumeth.2016.03.001

Marzolla, M. C. (2018). Tangled Up in Confounds—Unravelling the Controversial Roles of MTL-structures in Familiarity and Recollection. Maastricht Student Journal of Psychology and Neuroscience, 7, 7–20.

McDiarmid, A. D., Tullett, A. M., Whitt, C. M., Vazire, S., Smaldino, P. E., & Stephens, J. E. (2021). Psychologists update their beliefs about effect sizes after replication studies. Nature Human Behaviour, 5(12), 1663–1673. https://doi.org/10.1038/s41562-021-01220-7

O’Connor, A. R., Han, S., & Dobbins, I. G. (2010). The Inferior Parietal Lobule and Recognition Memory: Expectancy Violation or Successful Retrieval? The Journal of Neuroscience, 30(8), 2924–2934. https://doi.org/10.1523/JNEUROSCI.4225-09.2010

Ozubko, J. D., Moscovitch, M., & Winocur, G. (2017). The influence of recollection and familiarity in the formation and updating of associative representations. Learning & Memory, 24(7), 298–309. https://doi.org/10.1101/lm.045005.117

Peirce, J., Gray, J. R., Simpson, S., MacAskill, M., Höchenberger, R., Sogo, H., Kastman, E., & Lindeløv, J. K. (2019). PsychoPy2: Experiments in behavior made easy. Behavior Research Methods, 51(1), 195–203. https://doi.org/10.3758/s13428-018-01193-y

Ranganath, C., Yonelinas, A. P., Cohen, M. X., Dy, C. J., Tom, S. M., & D’Esposito, M. (2004). Dissociable correlates of recollection and familiarity within the medial temporal lobes. Neuropsychologia, 42(1), 2–13. https://doi.org/S0028393203001878 [pii]

Rohatgi, A. (2020). WebPlotDigitizer, Version 4.2 (4.2) [Computer software]. https://automeris.Io/WebPlotDigitizer.

Shrout, P. E., & Rodgers, J. L. (2018). Psychology, Science, and Knowledge Construction: Broadening Perspectives from the Replication Crisis. Annual Review of Psychology, 69(1), 487–510. https://doi.org/10.1146/annurev-psych-122216-011845

Simmons, J. P., Nelson, L. D., & Simonsohn, U. (2011). False-Positive Psychology: Undisclosed Flexibility in Data Collection and Analysis Allows Presenting Anything as Significant. Psychological Science, 22(11), 1359–1366. https://doi.org/10.1177/0956797611417632

Skavronskaya, L., Moyle, B., & Scott, N. (2020). The Experience of Novelty and the Novelty of Experience. Frontiers in Psychology, 11, 322. https://doi.org/10.3389/fpsyg.2020.00322

Szucs, D., & Ioannidis, J. P. (2020). Sample size evolution in neuroimaging research: An evaluation of highly-cited studies (1990-2012) and of latest practices (2017-2018) in high-impact journals. NeuroImage, 221, 117164. https://doi.org/10.1016/j.neuroimage.2020.117164

Tustison, N. J., Cook, P. A., Klein, A., Song, G., Das, S. R., Duda, J. T., Kandel, B. M., van Strien, N., Stone, J. R., Gee, J. C., & Avants, B. B. (2014). Large-scale evaluation of ANTs and FreeSurfer cortical thickness measurements. Neuroimage, 99, 166–179. https://doi.org/10.1016/j.neuroimage.2014.05.044

Wang, H., & Yushkevich, P. A. (2013). Multi-atlas segmentation with joint label fusion and corrective learning-an open source implementation. Front Neuroinform, 7, 27. https://doi.org/10.3389/fninf.2013.00027

Wilson, M. D. (1988). The MRC Psycholinguistic Database: Machine Readable Dictionary, Version 2. Behavioral Research Methods, Instruments and Computers, 20, 6–11.

Wixted, J. T., & Squire, L. R. (2011). The medial temporal lobe and the attributes of memory. Trends in Cognitive Sciences, 15(5), 210–217. https://doi.org/10.1016/j.tics.2011.03.005

Yonelinas, A. P. (1994). Receiver-operating characteristics in recognition memory: Evidence for a dual-process model. Journal of Experimental Psychology: Learning, Memory, and Cognition, 20(6), 1341–1354. https://doi.org/10.1037/0278-7393.20.6.1341

Yonelinas, A. P. (2001). Components of episodic memory: The contribution of recollection and familiarity. Philosophical Transactions of the Royal Society of London. Series B, 356(1413), 1363–1374. https://doi.org/10.1098/rstb.2001.0939

Yu, S. S., & Rugg, M. D. (2010). Dissociation of the electrophysiological correlates of familiarity strength and item repetition. Brain Research, 1320, 74–84. https://doi.org/10.1016/j.brainres.2009.12.071

